# Monitoring Gene Expression in Retina with synthetic serum markers

**DOI:** 10.64898/2026.01.16.699406

**Authors:** Jiaxiong Lu, Sangsin Lee, Meng Wang, Karen Zheng, Soo Hwan Oh, Jocelyn Lee, Sujung Soh, Yexuan Cao, Xuan Bao, Yifan Huang, Shirin Nouraein, Zigang Wu, Jerzy O. Szablowski, Rui Chen

## Abstract

Gene expression underlies retinal development, function, and pathogenesis. However, monitoring retinal gene expression in vivo is challenging. In this study, we apply synthetic serum markers, called Released Markers of Activity, or RMAs, to quantify gene expression in an intact retina through a simple blood test. We show that RMAs can quantify transduction of multiple retinal cell-types and monitor cell implantation or cell loss. We found RMAs are sensitive enough to measure transduction of rare cell populations, such as retinal ganglion cells and could detect transplantation of as few as 100 stem cells. Expression of RMAs showed no evidence of cell loss or immune activation in the retina. Overall, these findings demonstrate that RMAs are highly sensitive, tissue non-destructive gene expression reporters that can continuously track gene delivery and cell survival in intact retina through a simple blood test.

## INTRODUCTION

The ability of assessing cell activity, state, and function in living organisms is critical for us to better characterize cell function, diagnosing disease, and developing new therapeutic strategies^1^. Retina is one of the most accessible tissues in our body, and many noninvasive tools have been developed to benefit retinal research and clinical care^2^. For example, imaging methods such as optical coherence tomography (OCT)^3,4^, fundus photography^5^, and adaptive optics (AO)^6^ allow for detailed assessment of retinal histology. The electrophysiology and function of the retina can be evaluated with tools like electroretinography (ERG) and functional assessments^7^. Although these methods offer high-resolution analysis of retinal structures at the cellular level or assessment of function, their ability to measure gene expression is limited. One could theoretically use a fluorescently-labeled tag to monitor gene expression in the retina using fluorescent microscopy, but in practice, this approach uses strong light sources leading to cellular phototoxicity^8^, physical retinal damage^9–11^, and confounds the results through activation of photoreceptors^11^. Consequently, most research settings still rely on post-mortem histology and biochemical analysis, which prevents longitudinal studies. Another limitation of current methods is their low sensitivity in detecting and characterizing the functions of rare retinal cell populations. To address this challenge, it is essential to develop accessible, minimally invasive tools with high sensitivity and specificity that can detect molecular changes even within small subsets of retinal cells. Recently, a new class of genetically encodable reporters called Released Markers of Activity (RMAs)^12,13^ enabled monitoring of gene expression in deep tissues with a simple blood test. When expressed in the brain, RMAs enabled multiplexed, noninvasive, and site-specific monitoring of brain gene expression. In our original study, RMAs consisted of two protein domains – the naturally secreted *Gaussia* luciferase (Gluc) and the fragment crystallizable (Fc) region of an IgG2 antibody which allowed the RMAs to travel across an intact blood-brain barrier (BBB) through a process of reverse transcytosis^12–15^. RMAs were able to monitor gene delivery, allowing for the detection of gene expression in as few as ~12 neurons, and predicted the efficiency of gene delivery to the brain^12,13^. When expressed under a c-Fos responsive gene circuit, RMAs could also monitor chemogenetic activation of neuronal activity through a blood test^12^. We therefore hypothesized that RMAs could also be a valuable tool to quantify gene expression in the living retina.

The retinal vasculature shares a similar architecture with the brain and features the blood-retinal barrier (BRB) that isolates its cell populations from peripheral blood^16^. Similarly to the brain, intravitreally delivered antibodies are cleared from the retina through the neonatal Fc receptor (FcRN)^17–19^, which mediates reverse transcytosis by binding of the Fc^20^. Thus, we hypothesized that RMAs expressed in the retina could cross the BRB, enter the blood, and allow for sensitive, minimally invasive measurement of gene delivery, endogenous gene expression, or cell implantation and survival within the retina. RMAs’ sensitivity makes them particularly well-suited to track rare retinal cell types such as amacrine cells (ACs) or retinal ganglion cells (RGCs), which respectively account for about 8% and 2% of the retinal population and cannot be easily tracked through available methods^21–24^. Finally, RMAs, because of their non-tissue destructive readout, could measure the gene expression repeatedly in the same cell population, enabling longitudinal monitoring of gene expression within the same animal. Such longitudinal monitoring is critical to understanding the causality of gene expression on disease progression and cell survival^25^.

To validate whether RMAs could be adopted to monitor retinal physiology, we first expressed them in the common and rare cells in the retina through gene delivery of adeno-associated viral vectors (AAVs), finding significantly elevated levels of RMAs in the blood. When expressed within the stem cells, RMAs monitored the success of initial transplantation of stem cells and their subsequent survival. Furthermore, we have shown that RMAs can be used to quantify targeted cell population degeneration with high sensitivity and precision. Overall, this work demonstrates a new method for monitoring transgene expression within the retina using a simple blood test with versatile applications for both preclinical and clinical studies.

## RESULTS

### Detection of retinal transduction with a blood draw using RMAs

To test whether RMAs can cross the BRB into the blood, we delivered gene encoding RMA through a subretinal injection of AAV2. The AAV carried Gluc-based RMA (GlucRMA) driven by a constitutive EF1a promoter (Fig. 1a) which is known to drive gene expression across multiple cell populations of the retina^26,27^. Blood sample from each mouse was collected as the baseline before subretinal injection of AAV2 at a total dose of 6.24E9 viral particles (VP). Additional batches of blood samples were collected from each animal after 3 and 5 weeks post-injection and all blood samples were analyzed for RMA signal together using a luciferase assay as detailed in the Methods section. The RMA fold change was used as the primary readout, as it enables standardized comparisons across different instruments. Notably, we observed significant elevations in RMA levels in the blood serum, with over a 48,000-fold signal (*p*<*0.05*, one-way ANOVA with Tukey-HSD, n=5, Fig. 1b) compared to baseline, suggesting that the AAVs successfully transduced mouse retinal cells and released RMAs into the bloodstream. We did not observe differences in RMA levels between weeks 3 and 5 (P=0.33, one-way ANOVA with Tukey-HSD, n=5) (Fig. 1b). This finding suggests that blood RMA levels reached a steady state by the 3rd week after injection and remained stable until at least the 5th week. To identify the number of RMA-transduced cells, we collected the retinas at the 3rd and 5th week post-injection, and immunostained the retinas for Gluc and the photoreceptor and bipolar cell marker Otx2. We observed positive Gluc expression at the injection site (Fig. 1c), with 42.6% (SD±3.8, n=3) and 48.7% (SD±2.1, n=3) of Gluc and Otx2 double-positive cells being detected at the 3- and 5-week time points, respectively (Fig. 1d). Together, these results indicate that RMAs expressed in the retina can cross the BRB and become detectable in blood, enabling measurement of the retinal transgene expression with a blood test.

**Figure 1.**
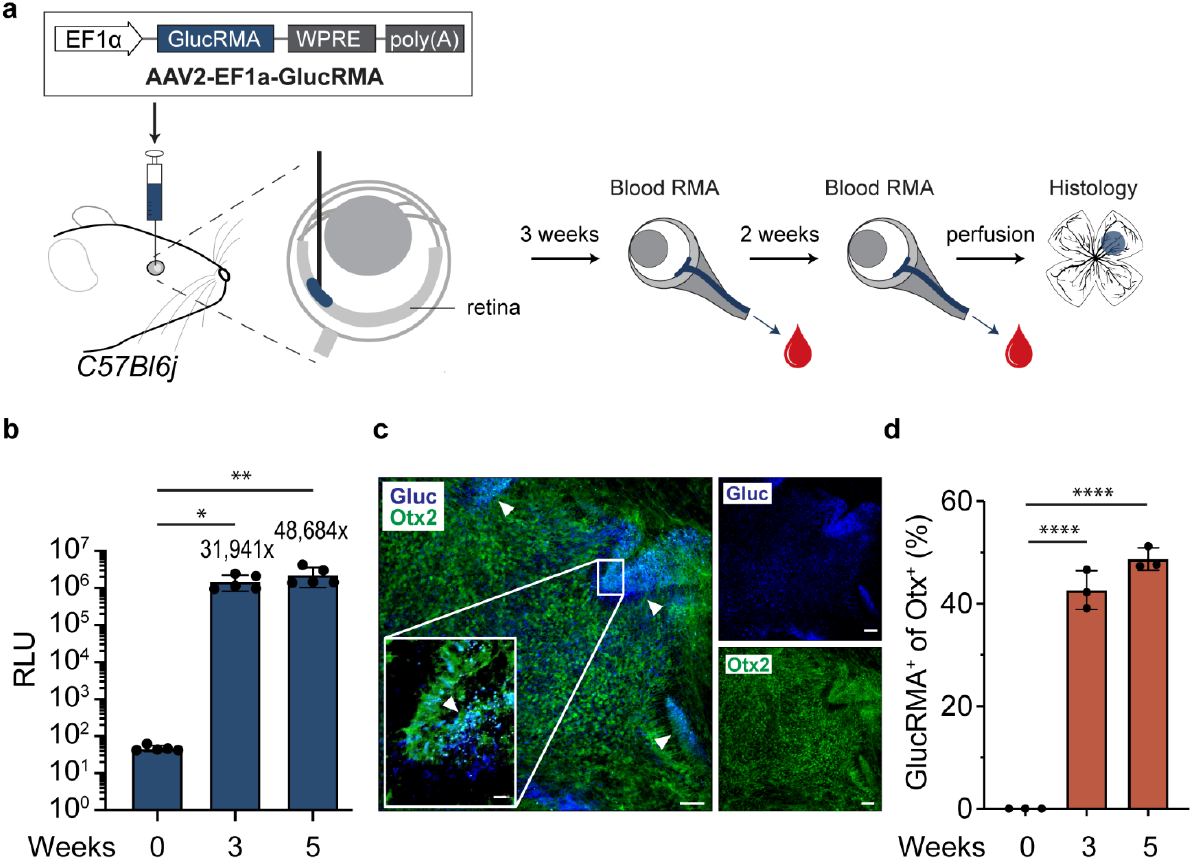
RMA enables efficient labeling and gene expression monitoring of retinal cells in mice. **a**. Schematic diagram depicting labeling of mouse retinal cells with GlucRMA followed by blood collection for RMA measurements and histology to confirm its retinal expression. **b**. Bioluminescence signals of GlucRMA in blood samples at 3 and 5 weeks post-AAV transduction. The numbers above bars indicate the signal fold changes relative to baseline at 0 weeks. n=5 mice analyzed. One-way ANOVA with Tukey’s test (F2,12=9.748, *P=0.0031*). *P=0.0362* (0 vs. 3 weeks), *P=0.0025* (0 vs. 5 weeks), *P=0.3278* (3 vs. 5 weeks). **c**. Whole mounts of injected retinas co-stained with antibodies against Gluc and Otx2, a marker specific to photoreceptor and bipolar cells. White arrows indicate regions where both markers are colocalized. Scale, 50 μm (main image) and 20 μm (insets). **d**. Quantification of GlucRMA-positive cells within the Otx-positive cells at 3 and 5 weeks post-AAV transduction. n=3 independent samples analyzed. One-way ANOVA with Tukey’s test (F2,6=335.2, *P*<*0.0001*). *P*<*0.0001* (0 vs. 3 weeks), *P*<*0.0001* (0 vs. 5 weeks), and *P=0.0572* (3 vs. 5 weeks). *\*P*<*0.05, **P*<*0.01, ****P*<*0.0001*. Data are shown as mean ± s.d.

### RMA tracks gene expression of a rare cell population in the retina

Given the presence of RMA in blood at over 48,000-fold, we hypothesized that it would be possible to detect RMA in blood even if only a small number of cells in the retina expressed RMA. To test this, we performed intravitreal injection of AAV2 carrying floxed GlucRMA reporter gene into Vglut2-cre mice eyes, which express Cre specifically in RGCs (Fig. 2). Blood samples were collected prior to the injection and then at 3 and 5 weeks after injection to quantify RMA levels using a luciferase assay. We observed a significant increase in RMA levels in serum at 3 and 5 weeks, showing over a 640-fold rise compared to the baseline (p<0.001, one-way ANOVA with Tukey-HSD, n=23, Fig. 2b). This fold change was lower than the one observed from labeling major retinal cell populations, which is consistent with the lower abundance of RGCs. To validate whether blood RMA levels correspond to the cell infection rate, we collected retinas at 3 and 5 weeks post-AAV2 injections, and immunostained them against Gluc and the RGC marker (Brn3a). We observed a significant increase in Gluc expression within RGCs (Fig. 2c), with 20.2%±3.85 (mean ± 95% CI, n=3) and 25.4%±6.2 (mean ± 95% CI, n=3) of cells being positive for both Gluc and Brn3a at 3 and 5 weeks, respectively, significantly higher than baseline controls (p<0.001, One-way ANOVA with Tukey HSD, N=3). These findings provide evidence that RMAs are released from the retina into the bloodstream and can monitor the transduction of a rare retinal cell type.

**Figure 2.**
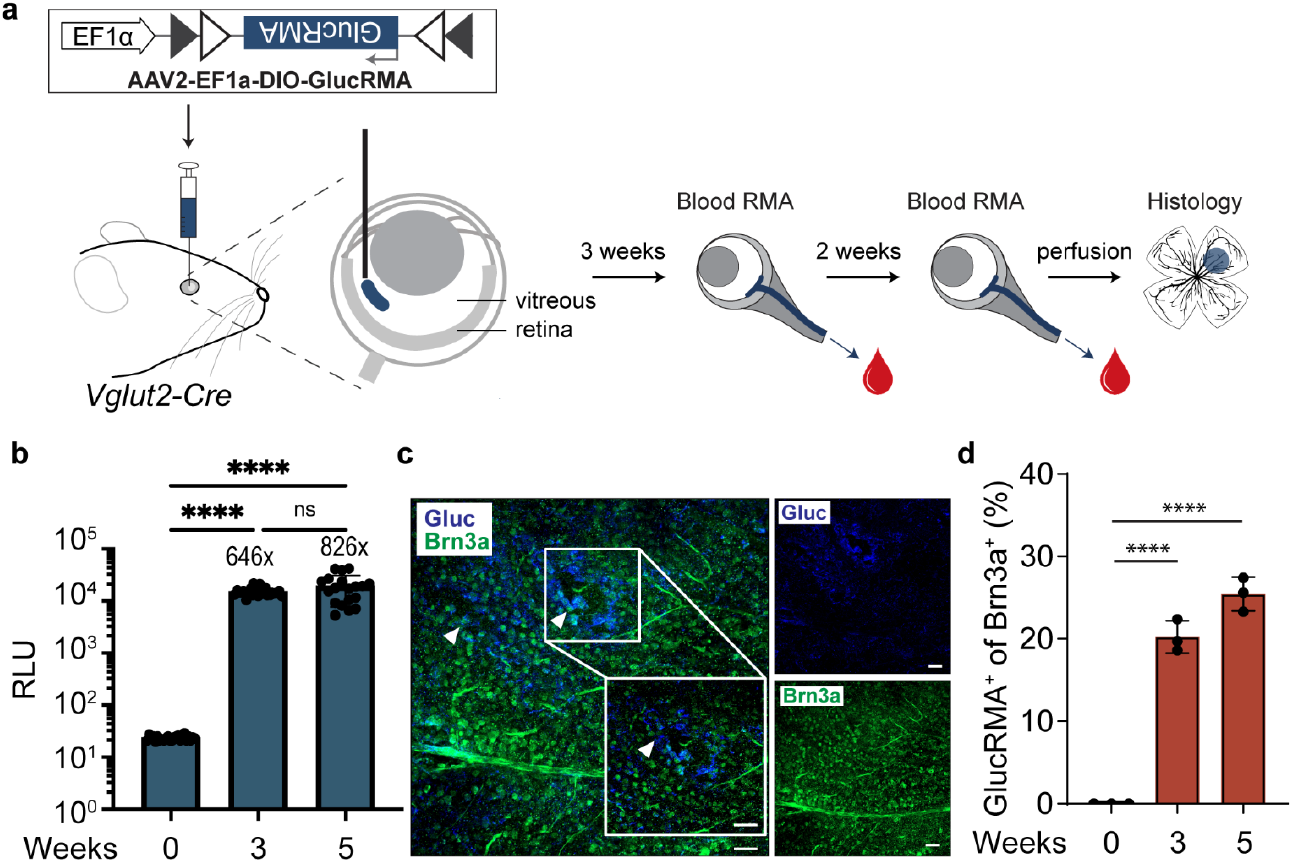
RMA allows selective measurement of gene expression in rare retinal cell types in mice. **a**. Schematic illustrating the selective labeling of rare RGCs in mouse retina. The GlucRMA cassette is positioned in a double-inverted orientation flanked by loxP and lox2272 sites, which undergo Cre-mediated recombination to revert to correct orientation and restore gene expression. **b**. Bioluminescence signals of GlucRMA detected in blood at 3 and 5 weeks after AAV transductions. Signal fold changes relative to baseline at 0 weeks are indicated above bars. n=24 mice analyzed. Unpaired t-test (t =27.5, df<45). *P*<*0.0001* (0 vs. 3 weeks). **c**. Whole mounts of injected retinas co-stained with antibodies against Gluc and Brn3a, a marker specific to RGCs. White arrows point to areas where both markers are colocalized. Scale, 50 μm (main image) and 20 μm (insets). **d**. Quantification of GlucRMA-positive cells among the Brn3a-positive cells at 3 and 5 weeks post-AAV transduction. n=3 independent samples analyzed. One-way ANOVA with Tukey’s test (F2,6=201.9, *P*<*0.0001*). *P*<*0.0001* (0 vs. 3 weeks), *P*<*0.0001* (0 vs. 5 weeks), and *P=0.0190* (3 vs. 5 weeks). *\*\*\*\*P*<*0.0001*. Data are shown as mean ± s.d.

### RMAs monitor photoreceptor cell transplantation in vivo

Photoreceptors or retinal pigment epithelium (RPE) cells^28,29^ replacement therapy represents one of the highly promising strategies for restoring vision in advanced retinal degenerative diseases^29–31^. Previous studies demonstrated that photoreceptor cells derived from ES and iPSCs can survive and integrate into the host retina following transplantation^32,33^, highlighting their potential as a therapeutic strategy. Notably, RPE replacement therapies are advancing in the clinic, with some nearing phase II clinical trials to determine efficacy in humans and progressing toward phase III and IV in several ongoing clinical trials^31,34^. The ability to measure transplantation efficiency and stability would be a valuable tool in studying and applying cell replacement therapies. Here, we decided to establish whether RMAs could monitor the retinal transplantation of the iPSC-derived photoreceptors and RPEs. To achieve this, we expressed RMAs in transplanted cells under constitutive cell-type specific promoters and used blood draws to monitor survival of transplanted cells in the retina. Three-month-old retinal organoids were used to generate differentiating photoreceptor cells for transplantation (see Materials and Methods). Lentiviral vector (Lenti-GRK-GlucRMA-GFP), which contains a photoreceptor-specific GRK promoter driving the expression of GlucRMA and GFP tag, transduced 3-month-old retinal organoids at a dose of 5E+08 VP per organoid (5e4 cells/retina organoid) (Fig 3a). Fourteen days post-infection, the retinal organoids were dissociated into single cells and subjected to FACS sorting to isolate GFP-positive (1.7%, Fig. S2) transduced cells. We then transplanted 10,000 FACS-sorted GFP+ photoreceptors into each SCID mouse eye via subretinal injections. Subsequently, we collected blood samples at 4- and 8-week post-transplantation to measure serum RMA levels compared to the baseline. Our results demonstrated a significant, 439±19 (mean ± 95% CI)-fold, elevations of RMA signals in blood at both time points compared to the baseline controls (n=5, p<0.0001, One-way ANOVA with Tukey-HSD, for 4-week and 8-week time points). In parallel, we performed immunofluorescence (IF) against Gluc, GFP, and Ku80 (a human cell-specific antigen) to detect transplanted cells in the mouse retinas. We observed co-localization of these markers, indicating successful transplantation of photoreceptors into the mouse retinas, consistent with the release of RMAs detected in the blood samples (Fig 3c, Fig. S3a). Furthermore, IF analysis of Gluc, GFP, and Ku80 of the cross-sections of injected retinas showed that the transplanted photoreceptors were located in the ONL or between the outer nuclear layer (ONL) and inner nuclear layer (INL) (Fig. S3b to d). A small number of cells, 112±16 cells (mean ± 95% S.D., n=5), were detected in each retina, suggesting high sensitivity of RMA (Fig. S3e).

**Figure 3.**
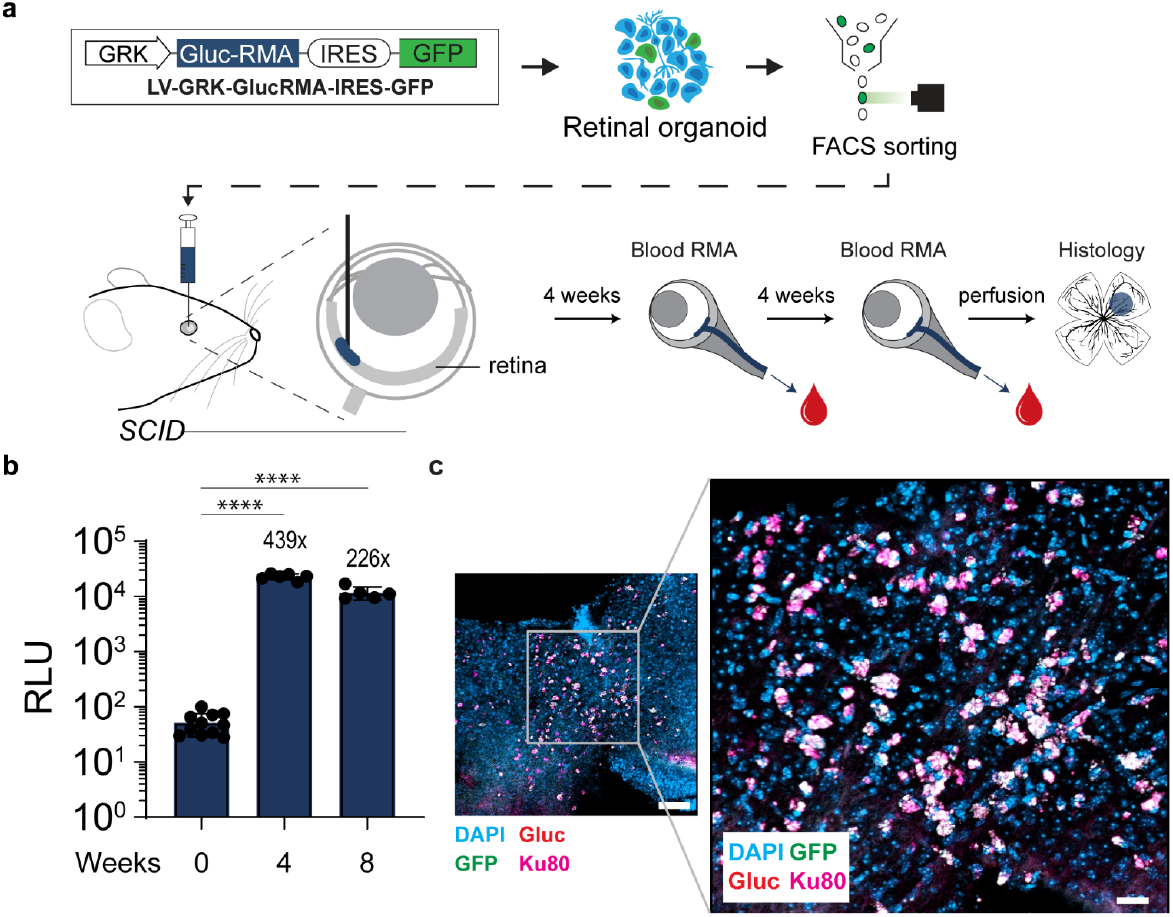
Noninvasive monitoring of photoreceptor cells transplantation in retina using RMAs. **a**. Schematic illustrating the procedure for monitoring photoreceptor cell transplantation. Lentiviral vectors encoding GlucRMA and GFP were transduced into retinal organoids, and FACS-isolated GFP-positive cells were transplanted into the retina of SCID mice, followed by blood collection to monitor successful integration. **b**. Bioluminescence signals of GlucRMA measured in blood samples at 4 and 8 weeks after lenti-GRK-RMA-GFP transduction. Signal fold changes relative to baseline at 0 weeks are indicated above bars. n=11 (baseline), 6 (4 weeks), and 5 (8 weeks) mice analyzed. One-way ANOVA (F2,19=248.1, *P*<*0.0001*). *P*<*0.0001* (0 vs. 4 weeks), *P*<*0.0001* (0 vs. 8 weeks), *P*<*0.0001* (4 vs. 8 weeks). **c and d**. Immunostaining of injected retinal whole mounts using anti-Gluc, anti-GFP, and anti-Ku80 antibodies. Scale, 50 μm (main image) and 20 μm (insets). Data are shown as mean ± s.d.

### Monitor RPE cell transplantation in vivo through RMAs

Similarly, to test RPE cell replacement, AAV2-EFS-GlucRMA-GFP transduced H9 stem cell-differentiated RPEs at a dose of 8.6E10 VP were subretinally injected into SCID mice (Fig 4a). We tested four different transplantation doses: 100,000, 10,000, 1,000, and 100 cells. Four weeks after transplantation, our results showed significant increases in RMA signals in the blood compared to baseline controls, with average signals of 341-, 142-, 38-, and 9-fold over the baseline with all cell doses showing a significant signal elevation (Q<0.000001, Q<0.000001, Q=0.000050, Q=0.000852, for each respective dose. Multiple t-test comparison with 5% FDR correction using Benjamini, Krieger, and Yekutieli method, Fig 4b, Fig. S4a to c). We also counted the transplanted and surviving RPE cells in dissected retinal whole mounts, observing an average of 16±5.1, 95.4±5.0, 405.4±55.3, and 5027.8±404.7 RPE cells for each corresponding dose (mean ± S.D., Fig. S4d). We observed that the blood RMA levels highly correlated with the number of transplanted cells as counted in post-mortem histology (R=0.988, P<0.0001) and similarly we observed a positive correlation between the injected cell dose and the number of surviving cells in the retina (R=0.99, P<0.0001, Fig. 4c).

**Figure 4.**
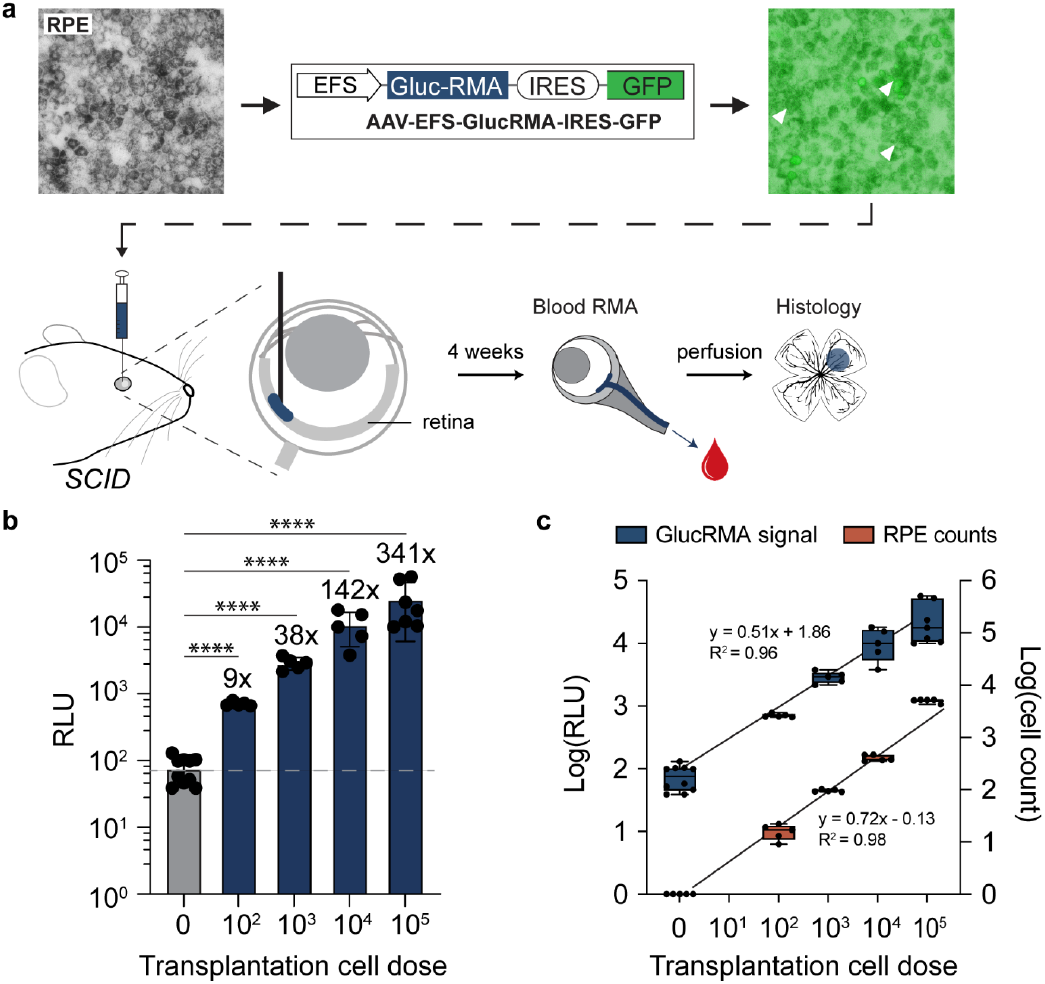
RMA allows for noninvasive monitoring of RPE cell transplantation. **a**. Schematic illustrating the process for monitoring RPE cell transplantation. **b**. Bioluminescence signals of GlucRMA measured in blood at 4 weeks after AAV-EFS-RMA-GFP-transduced RPE transplantation with varying doses of transplanted RPE cells. Signal fold differences compared to untransplanted control are shown above bars (9x-341x). n=10 (no transplantation), n=7 (102 RPEs), and n=5 (103, 104, and 105 RPEs) mice analyzed. (*Q*<*0.000001, Q*<*0.000001, Q=0.000050, Q=0.000852*, for each respective dose. Multiple t-test comparisons with 5% FDR correction using Benjamini, Krieger, and Yekutieli method). **c**. Pearson’s correlation between the transplantation dose and both blood RMA bioluminescence signals and the number of surviving RPE cells. *\*\*\*\*P*<*0.0001*. Data are shown as mean ± s.d.

### Tracking NMDA-induced cell death with RMAs

Our previous experiments demonstrated that RMA is a sensitive and reliable tool for monitoring transgene expression in the retina. To further demonstrate the potential utility of RMA, we tested whether RMAs could monitor degeneration of targeted rare cell types. As described above, we specifically labeled RGCs with RMA by injecting AAV-DIO-RMA into Vglut2-Cre mice. Three weeks after the AAV injection, the mice were treated with NMDA, which induces RGC cell death^35,36^. Indeed, 4 weeks after NMDA treatment, we observed a significant reduction in the number of RGCs (Fig. 5a, Fig. S5a and b). Blood RMA levels were collected from four groups 4 weeks after NMDA treatment: 1) untreated baseline control, 2) AAV-DIO-RMA-injected only with NMDA treatment, 3) AAV-DIO-RMA + NMDA-treated, and 4) NMDA treatment only mice. The results showed that, compared to baseline controls, the blood RMA levels were elevated by 284.2±170.7-fold in the AAV-DIO-RMA group, 10.9±5.5-fold in the AAV-DIO-RMA + NMDA group, and 2.0±1.3-fold in the NMDA-only group (mean ± S.D.; Q<0.0001, Q<0.0001, and Q>0.05, respectively, Multiple t-test comparison with 5% FDR correction using Benjamini, Krieger, and Yekutieli method, Fig. 5b). As expected, no significant RMA change was observed between the baseline and NMDA treatment only group (2-paired t-Test, P>0.9999). We did observe a 284-fold increase of RMA level in the AAV-DIO-RMA group compared to the baseline. Notably, the combination of AAV-DIO-RMA and NMDA treatment resulted in a 96.2±3.0% (mean ± S.D.) reduction in blood RMA levels compared to AAV-DIO-RMA alone, consistent with the RGC degeneration upon NMDA treatment (Fig. 5b). Indeed, we observed a strong positive correlation between the cell count and blood Gluc-RMA signal for the GlucRMA+; NMDA+ group in post-mortem histology (Pearson correlation coefficient, R = 0.97, P =0.0042, Fig. 5c). Additionally, AAV-DIO-RMA-injected retinas exhibited minimal activation of Müller glial cells, as indicated by low GFAP expression, and minimal microglial activation, as shown by limited Iba1 staining (Fig. 5d). Importantly, RGC cell death, marked by reduced RBPMS expression, was observed only in NMDA-treated retinas and not in AAV-DIO-RMA-injected retinas (Fig. 5d).

**Figure 5.**
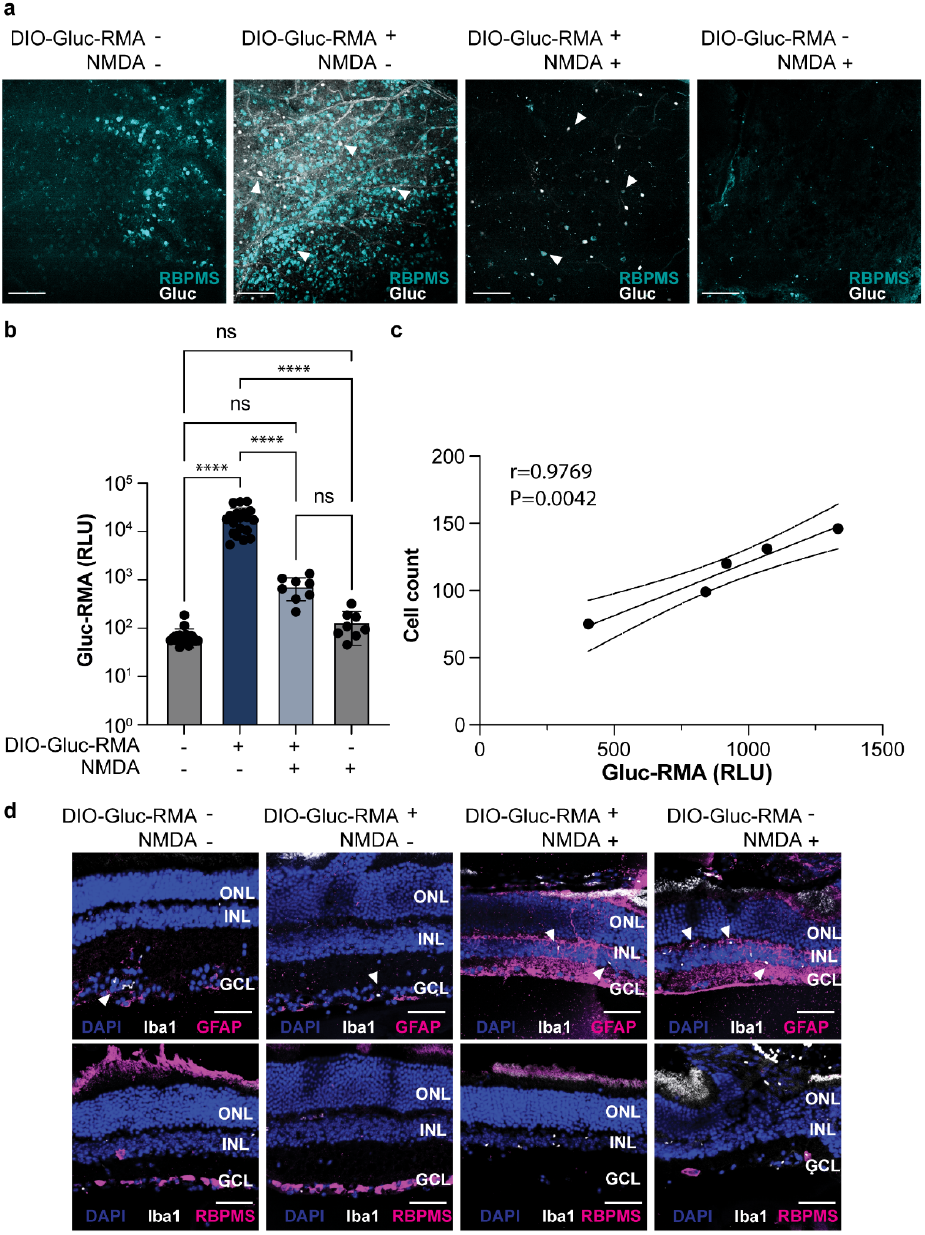
RMA-Based Monitoring of NMDA-Induced Cell Death. **a**. Representative images of RBPMS and Gluc immunostaining in retinal whole mounts from mice injected with or without AAV-DIO-RMA and NMDA. Scale bar = 100 μm. **b**. Bioluminescence signals of GlucRMA measured in blood at 4 weeks post-injection of AAV-DIO-RMA with or without NMDA. Signal fold changes compared to baseline controls: AAV-DIO-RMA group, 284.2±170.7-fold; AAV-DIO-RMA + NMDA group, 10.9±5.5-fold; NMDA-only group, 2.0±1.3-fold (mean ± S.D.; *Q*<*0.000001, Q=0.9945, Q*>*0.9999*). Multiple t-tests with 5% FDR correction using Benjamini, Krieger, and Yekutieli method. **c**. Correlation between the cell count and Gluc-RMA signal for the GlucRMA+; NMDA+ group. *\*\*P*<*0.0042*. Data presented as mean ± s.d. **d**. Representative images of Iba1, GFAP, and RBPMS immunostaining on retinal paraffin sections from mice injected with or without AAV-DIO-RMA and NMDA. Scale bar = 50 μm.

## DISCUSSION

RMA technology represents a significant advancement in retinal monitoring, providing a minimally invasive approach to evaluate retinal health and cellular function. In contrast to traditional techniques such as ERG, OCT, and AO, which primarily assess retinal structure and overall functional output, RMA offers complementary insights by directly monitoring gene expression within retinal cell populations. Because RMA measures gene expression dynamics, it holds distinct advantages for studying rare retinal cell types. Monitoring gene expression in these cells is particularly important, as current in vivo monitoring systems often suffer from limited sensitivity and rely on complex imaging procedures that can introduce phototoxic effects. Furthermore, existing methods can confirm the survival and localization of transplanted cells but cannot determine their functional integration. For example, global functional assays like ERG frequently fail to differentiate the specific contributions of transplanted cells before and after transplantation^37^.

In this study, we successfully expressed the RMAs under non-selective constitutive promoters following subretinal/intravitreal injection. We monitored two distinct types of retinal cells: photoreceptors, which are the predominant cell type in the retina, and RGCs, which constitute approximately 1% of the retinal cells^38^. In both cases, RMA expressions were sufficient to enable detection with 5 µl of plasma (Fig. 1b and Fig. 2b). Building on this success, we sought to explore the potential utility of RMAs in monitoring cell therapy. Human embryonic stem cell-derived RO and RPE cells have shown promise as sources of cells for transplantation^39–43^. Our results suggested that both RO (Figure 3c) and RPE (Figure 4c) successfully integrated into the retina and continued to express RMAs for at least 8 weeks (Figure 3b). This result is of particular interest for clinical applications, as currently there are no good methods to monitor the numbers of transplanted cells and their survival in the retina.

It is important to note that although the BBB shares many structural and functional similarities with the inner BRB, these similarities do not apply to the choroidal circulation, which supplies the outer retina. The outer retina is the primary site for most retinal gene therapies over the past two decades and is also where cellular replacements are initially targeted^44,45^. In contrast, the inner and mid-retina are compartmentalized away from the outer retina. In this study, we used both subretinal and intravitreal delivery paradigms to target the inner and mid-retina. All our current results support the conclusion that RMA is produced and secreted by retinal cells and subsequently diffuses to nearby capillaries. This is further supported by our subretinal injection of purified RMA protein, which led to over a 10,000-fold increase in blood RMA levels (Fig. S6), demonstrating that RMA proteins can traverse retinal tissue and enter the circulation. Nevertheless, the precise pathway by which RMA proteins penetrate the retina remains to be elucidated.

Cell-based therapies are often genetically modified^46,47^, including clinically approved treatments such as Abecma (idecabtagene vicleucel), which is the first cell-based gene therapy approved for multiple myeloma^48^. This therapy involves genetically engineering a patient’s T-cells to specifically target and destroy myeloma cells. Another example is afamitresgene autoleucel (Tecelra), which is used to treat certain cases of metastatic synovial sarcoma^49^. By introducing RMA or similar construct into the cell therapies one could monitor the success of therapy to provide information on the therapeutic efficacy and the need for re-administration. Further, by placing RMAs under the control of endogenous cell state-dependent promoters^50,51^ it may be possible to glean further information about the functionality of these cell transplants over time.

To test if RMAs could be used to monitor targeted cell population degeneration, in addition to implantation, we used an NMDA induced RGC degeneration model^36,52^. We observed a 96.2% reduction in blood RMA levels four weeks after NMDA injection, demonstrating the capability of RMAs to monitor cell loss over time. Furthermore, AAV-delivered RMAs were found to be safe, as there was no significant neuronal loss and minimal activation of Müller glial cells or microglial cells (Fig. 5d). This indicates that AAV-RMA delivery does not elicit harmful immune responses in the retina, further supporting the safety of our system.

In this study, we chose SCID mice for cell transplantation due to their immunodeficient status, which minimizes the risk of immune rejection of transplanted human cells^51^. In humans, cell transplants are typically autologous, reducing the likelihood of immune responses. However, murine models of retinal degeneration are often immunocompetent. Therefore, to effectively evaluate the efficacy of cell transplants and the utility of RMAs in assessing these cells in retinal degeneration, the use of immunosuppressive drugs will be necessary. Compounds such as Cyclosporine^53,54^ and Tacrolimus (FK506)^55^ could be administered to mitigate immune responses and enhance the success of cell transplantation in more immunocompetent mouse models.

Future studies will include translation of RMAs for retinal monitoring of gene expression and cell transplants to large animals and possibly the clinic. Before that can happen, long-term studies of safety and efficacy will need to be performed. We recently took the first step towards translation, showing that RMAs can work in other species. By fusing the Fc region of a rhesus macaque IgG antibody to GLuc, forming NHP-RMA, we have shown successful BBB passage of NHP-RMA proteins and their ability to monitoring gene delivery to the brain^56^. To translate RMAs for the clinic, we will likely use human-derived Fc for endothelial transcytosis and use human-derived or humanized proteins for detection in the serum to avoid RMAs’ potential immunogenicity.

In summary, we show that RMA technology can monitor gene and cell delivery in various retinal cell populations. This technology enables minimal-invasive tracking of cellular and genetic activities within the retina following intraocular gene or cell delivery. When combined with gene and cell therapy, RMA technology could serve as a versatile tool that can be applied to both preclinical and clinical studies.

## MATERIALS AND METHODS

### Animal subjects

Male and female WT (Strain #000664) and Vglut2-Cre (Strain #028863) mice 57,58, aged 8-10 weeks, were purchased from the Jackson Laboratory. These mice were of the C57BL/6J genetic background and were used for AAV-based photoreceptor and RGC labeling. Cell transplantations (Photoreceptor and RPE) were performed on SCID mice. SCID mice had a genetic autosomal recessive mutation known as Prkdcscid. These SCID mice, homozygous for the Prkdcscid allele, exhibit severe combined immunodeficiency affecting both B and T lymphocytes, with the mutation mapped to the centromeric end of chromosome 16^59–61^. Both male and female mice were used in the study and were selected randomly. We have not observed differences between these groups of mice. All animals in this study were housed under a 14-hour light/10-hour dark cycle. The Institutional Animal Care and Use Committee (IACUC) at University of California, Irvine approved all animal procedures (Protocol AUP-24-092).

### Plasmid construction

All plasmid maps and their applications are provided in Fig S1. The constructed plasmids were based on templates from our previous study: pAAV-hSyn-GlucRMA-IRES-EGFP (Addgene #189629) and pAAV-hSyn-DIO-GlucRMA-IRES-EGFP (Addgene #189630). The EFS promoter, a short, modified version of the EF1α promoter, was used to construct pAAV-EFS-GlucRMA-IRES-EGFP. The hSyn promoter from pAAV-hSyn-GlucRMA-IRES-EGFP was removed by digesting the plasmid with XbaI. The EFS promoter was amplified from pAAV-EFS-SpCas9 (Addgene #104588) and extracted using the QIAquick Gel Extraction Kit (Qiagen). The digested backbone and the EFS segment were then assembled using Gibson Assembly. To construct pAAV-EFS-DIO-GlucRMA-IRES-EGFP, we again removed the hSyn promoter from pAAV-hSyn-DIO-GlucRMA-IRES-EGFP using XbaI digestion and inserted the EFS promoter through Gibson Assembly.

To generate Lenti-GRK-GlucRMA-GFP, plasmid pXPR_011 (Plasmid #59702) was digested using XmaI and MluI, followed by purification of the 6,452 bp vector with the QIAquick Gel Extraction Kit (Qiagen). Concurrently, PCR was performed to create inserts containing homology arms of hGRK1, specifically a 292 nt segment of the human G-protein-coupled receptor protein kinase 1 promoter (positions 1793 to 2087, GenBank AY327580), and the GlucRMA-IRES-EGFP sequence sourced from pAAV-hSyn-GlucRMA-IRES-EGFP (Addgene #189629). The PCR was conducted using Q5® High-Fidelity DNA Polymerase (NEB #M0491L). Finally, Gibson Assembly was employed with NEBuilder® HiFi DNA Assembly Master Mix (NEB #E2621L) to ligate the digested vector with the GRK1 promoter and GlucRMA-IRES-EGFP.

The plasmid constructs have been deposited in Addgene. Preparation of live single-cell suspensions from and cultured H9-RPE and retina organoid

AAV-transduced retina organoids were harvested. Each retina organoid underwent dissociation using papain-based enzymatic digestion, following established procedures with slight modifications^62^. Specifically, 45 U of activated papain solution (Worthington, Cat. #LS003126), along with 1.2 mg L-cysteine (Sigma) and 1200 U of DNase I (Affymetrix) in 5 ml of HBSS buffer, was added to the retina organoid, and the mixture was incubated at 37 °C for 15 minutes to release live cells. Following incubation, the single-cell solution underwent centrifugation at 200 x g, and the papain was deactivated using ovomucoid solution (15 mg ovomucoid from Worthington Biochemical and 15 mg BSA from Thermo Fisher Scientific in 10 ml of MEM from Thermo Fisher Scientific).

The remaining retina organoids were triturated in an additional ovomucoid solution and filtered through a 20 nm plastic mesh. This process was iterated until the entire retina was dissociated. The collected cells, exhibiting over 90% viability, were stained with the Ready Probes cell viability imaging kit (blue/red) containing Hoechst 33342 and propidium iodide (R37610, Thermo Fisher Scientific). For capturing, retinal cells were diluted to 3E4/ml with 1× PBS (Thermo Fisher Scientific), RNase inhibitor (NEB, 40 KU/ml), and Cell Diluent Buffer (Takara Bio #640167).

### Fluorescence-activated cell sorting (FACS)

FACS was conducted at the Flow Cytometry Core, University of California, Irvine. In the process, each dissociated cell underwent laser interrogation. The flow cytometer machine was set up to ensure that each individual cell entered a unique droplet upon exiting the nozzle tip. This droplet received an electronic charge determined by the cell’s fluorescence. Deflection plates manipulated the cells, guiding them into designated collection tubes through attraction or repulsion. Following this, the sorted GFP+ live cell populations were scrutinized to confirm the effectiveness of the cell sorting process.

### Subretinal and intravitreal injection

Postnatal day 21 (P21) mice were anesthetized via intraperitoneal injection with a combination drug, Rodent-III, consisting of ketamine (22 mg/kg), xylazine (4.4 mg/kg), and acepromazine (0.37 mg/kg). To dilate the pupils, tropicamide (1.0%; Generic #1168129) and phenylephrine (2.5%; Bausch Lomb Americas INC #42702010210) eye drops were applied. The cornea was then anesthetized using proparacaine hydrochloride (0.5%; Generic #2963726). A beveled 30-gauge needle (BD #305106 New Jersey) was used to make a puncture towards the caudal side of the eye between the sclera and cornea. Subsequently, a 32-gauge blunt syringe (Hamilton #418382 Nevada USA) was inserted into the vitreous cavity through the puncture hole and advanced until the needle tip was positioned between the retina and RPE (subretinal) or in the middle of the vitreous (intravitreal). Finally, 1-2 μL of AAV, cell suspensions, or NMDA (#m3262 Sigma) were injected into the mouse eyes.

### Retro-orbital blood collection

Mice were anesthetized with 1.5%-2% isoflurane in air or O_2_. Subsequently, 1-2 drops of 0.5% ophthalmic proparacaine were applied topically to the cornea of each eye. A heparin-coated microhematocrit capillary tube (Fisher Scientific, Catalog No. 22-362566) was placed into the medial canthus of the eye to puncture the retro-orbital plexus and withdraw 50-100 μl of blood. The collected blood was centrifuged at 1,500g for 5 minutes to isolate plasma, which was then stored at –20°C until use.

### Luciferase Assay

For Gluc substrate, 0.5 mM native coelenterazine (CTZ) stock (Nanolight Technology, 303) was dissolved in luciferase assay buffer (10 mM Tris, 1 mM EDTA, 1.2 M NaCl, pH 8.0) containing 66% DMSO and stored at −80 °C. Before measuring bioluminescence, the CTZ stock was diluted to 20 µM in luciferase assay buffer and kept in the dark at room temperature for 1 h. The luciferase assay was performed using Infinite M Plex microplate reader (Tecan) equipped with i-control software for measurements. 5 μl of plasma was mixed with 45 μl of PBS containing 0.001% Tween-20 in a black 96-well plate. The bioluminescence of GlucRMA was measured using the microplate reader by injecting 50 μl of 20 μM CTZ dissolved in luciferase assay buffer into the plasma sample.

### Immunostaining

Immunofluorescent staining on cryosections or paraffin sections was carried out following established protocols^63–65^. Enucleated eyes were embedded in OCT compound (SAKURA #4583) and stored at −80°C for at least 2 hours. Cryosections were cut at a thickness of 10 µm and washed with PBS. For paraffin sections, enucleated mouse eyes were fixed overnight at 4°C in freshly prepared Davidson’s fixative (40% formaldehyde, 35% ethanol, 10% acetic acid, and 53% H_2_O). After fixation, the eyes were washed twice with PBS buffer and dehydrated through a graded ethanol series (50%, 70%, 95%, and 100%). All washing and dehydration steps were carried out at 4°C with gentle shaking. The dehydrated eyes were cleared in xylene twice for 1 hour each at room temperature in a fume hood. They were then transferred to a pre-warmed 50% xylene/50% paraffin mix at 60°C for 1 hour, followed by overnight incubation in 100% paraffin at 60°C. The next day, the eyes were embedded in paraffin, and the paraffin-embedded tissue blocks were sectioned at a thickness of 7 µm. Subsequently, the slides were incubated for 1 hour in hybridization buffer (10% normal goat serum, 0.1% Triton X-100, in PBS) and left overnight with primary antibodies.

Primary antibodies used in this study include rabbit anti-Gluc (1:1,500, Nanolight Technology), anti-OTX2 (1:200, Fisher Scientific#BAF1979), anti-Brn3a (1:50, Santa Cruz Biotechnology#sc-8429; avoid cross-reactivity with Gluc antibodies), anti-XRCC5/Ku80 Antibody (1:500, Sigma# ZRB2058), anti-GFP (1:400, Rockland#600-101-215), anti-Iba1 (1:400 Abcam#107159), anti-GFAP (1: 200, Sigma #HPA056030), and anti-RBPMS (1:400, Novus #NBP2-20112). The following day, the slides were washed in PBS, incubated with secondary antibodies for 2 hours, followed by incubation with DAPI at room temperature. Finally, slides were mounted with anti-fade medium (Prolong; Invitrogen) and coverslipped. Fluorescent images were captured using a Zeiss Apotome.2 microscope (Zeiss Axio Imager), and ImageJ software was employed to count the number of transplanted cells. All immunostaining experiments were independently performed with three biological replicates.

### Statistical analysis

A two-tailed t-test with unequal variance was employed to compare two data sets. For comparisons between three data sets, one-way ANOVA with Tukey’s honestly significant difference post hoc test was utilized. Data sets involving two or more variables were compared using two-way ANOVA with Sidak’s multiple comparison tests. For larger numbers of data sets, 4 or more, Multiple t-test comparison with 5% FDR correction using Benjamini, Krieger, and Yekutieli method was used. Linear regression was applied to determine the correlation between the plasma signal and the number of transduced neurons. All P values were calculated using Prism (GraphPad Software), with statistical significance denoted as follows: ns (not significant), *\*P* < *0.05, **P* < *0.01, ***P* < *0.001, ****P* < *0.0001*.

### Data availability

The authors state that all data supporting the results of this study are included within the paper and its Supplementary Information. The raw and analyzed datasets can be obtained from the corresponding author upon reasonable request. The plasmids designed in this study will be available in Addgene.

## ACKNOWLEDGEMENTS

This research was funded by the Foundation Fighting Blindness (BR-GE-0613–0618-BCM) and the National Eye Institute (R01EY022356, R01EY020540, R01EY018571) (R.C.). Additional support was provided by the David and Lucile Packard Foundation (2021-73005) and the National Institute of Biomedical Imaging and Bioengineering, including the Trailblazer Award (R21EB033059) and DP2 Award (DP2EB035905) (J.O.S.).

## AUTHOR CONTRIBUTIONS

J.O.S. and R.C. conceived the project. J.L., S.L., J.O.S., and R.C. collaboratively designed the experiments. J.L. conducted plasmid construction, retinal injections, cell transplantation, FACS, immunostaining, imaging, and histological analyses. S.L. contributed to plasmid design and construction, blood collection, luciferase assays, and statistical analyses. M.W., K.Z., S.O., and J. Lee performed data analysis, cell quantification, plasmid cloning, retro-orbital blood collection. S.S., Y.X., and X.B. prepared retina organoids and differentiated H9-derived RPE cells. Y.H., Z.W., and S.N. carried out luciferase assays on collected blood samples. The manuscript was jointly written by J.L., S.L., J.O.S., and R.C., with input and edits from all contributing authors

