## Supplementary Figures for "Monitoring Gene Expression in Retina with synthetic serum markers"

| Vector Name | Vector size | Virus | Infected models |
|---|---|---|---|
| AAV-EF1a-GlucRMA | 6819 bp | AAV2 | *C57BL6J* |
| AAV-EF1a-DIO-GlucRMA | 6531 bp | AAV2 | *Vglut2-Cre* |
| Lenti-GRK-RMA-GFP | 9595 bp | Lentivirus | H9-retina organoids |
| AAV-EFS-RMA-GFP | 6911 bp | AAV2 | H9-RPE, HEK293T |

**Figure S1. Vector information for RMA constructs.** Tables for RMA constructs used in this study.

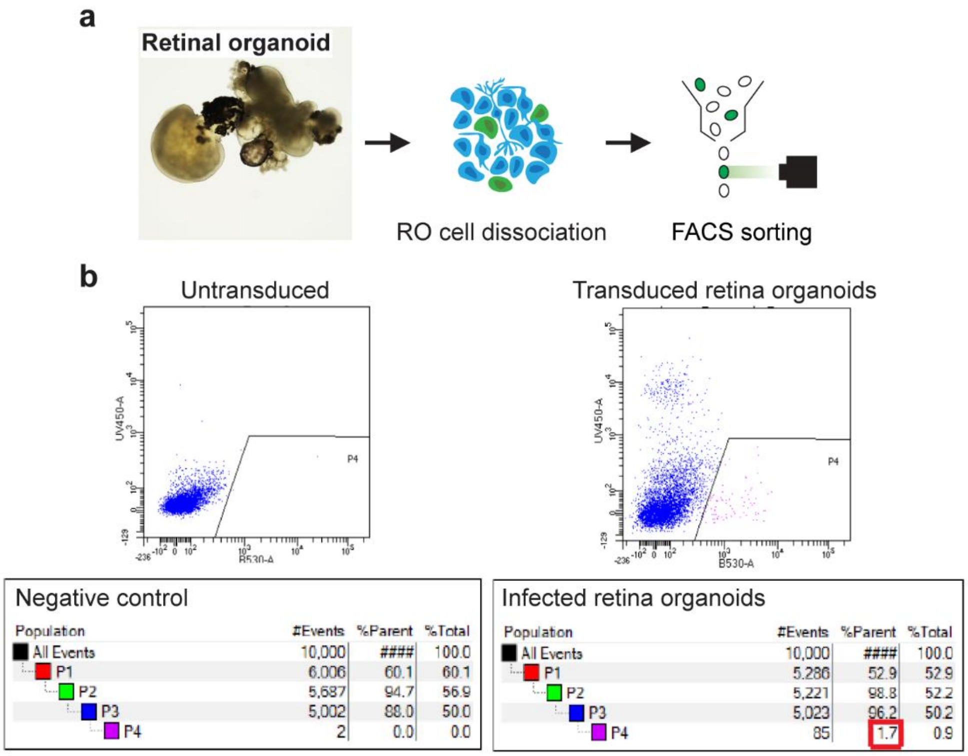


**Figure S2. Retina organoids as the source for photoreceptors. a.** Schematic illustrating the generate photoreceptors from cultured retina organoids. **b.** FACS sorting for GFP+ photoreceptors.

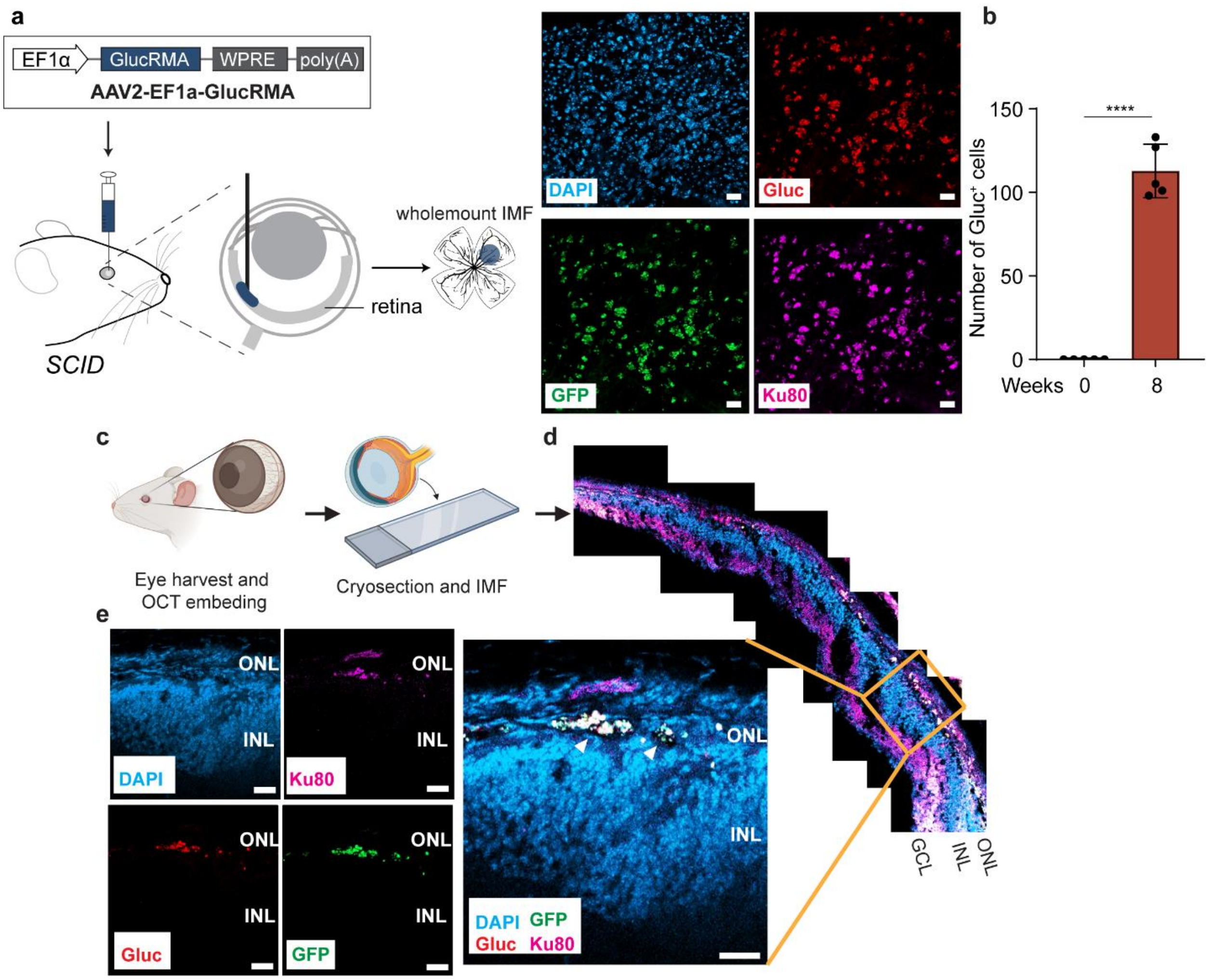


**Figure S3. Transplanted photoreceptors successfully integrate into host mouse retina. a.** Schematic illustrating immunostaining on retinal whole-mounts. **b.** Number of integrated Gluc$^+$ photoreceptors at 8 weeks post-transplantation. *n*=5 mice analyzed. Unpaired *t-test* (*t*=15.73, *df*=8). *P*<0.0001 (0 vs. 8 weeks) **c., d., and e.** Immunostaining of anti-Ku80, anti-Gluc, and anti-GFP and their localization to the ONL layer on photoreceptor transplanted retina cross sections. ONL, outer nuclear layer. INL, inner nuclear layer. GCL, ganglion cell layer. Scale, 20 µm (single channel), 20 µm (merged). ****$P$<0.0001. Data are shown as mean ± s.d.

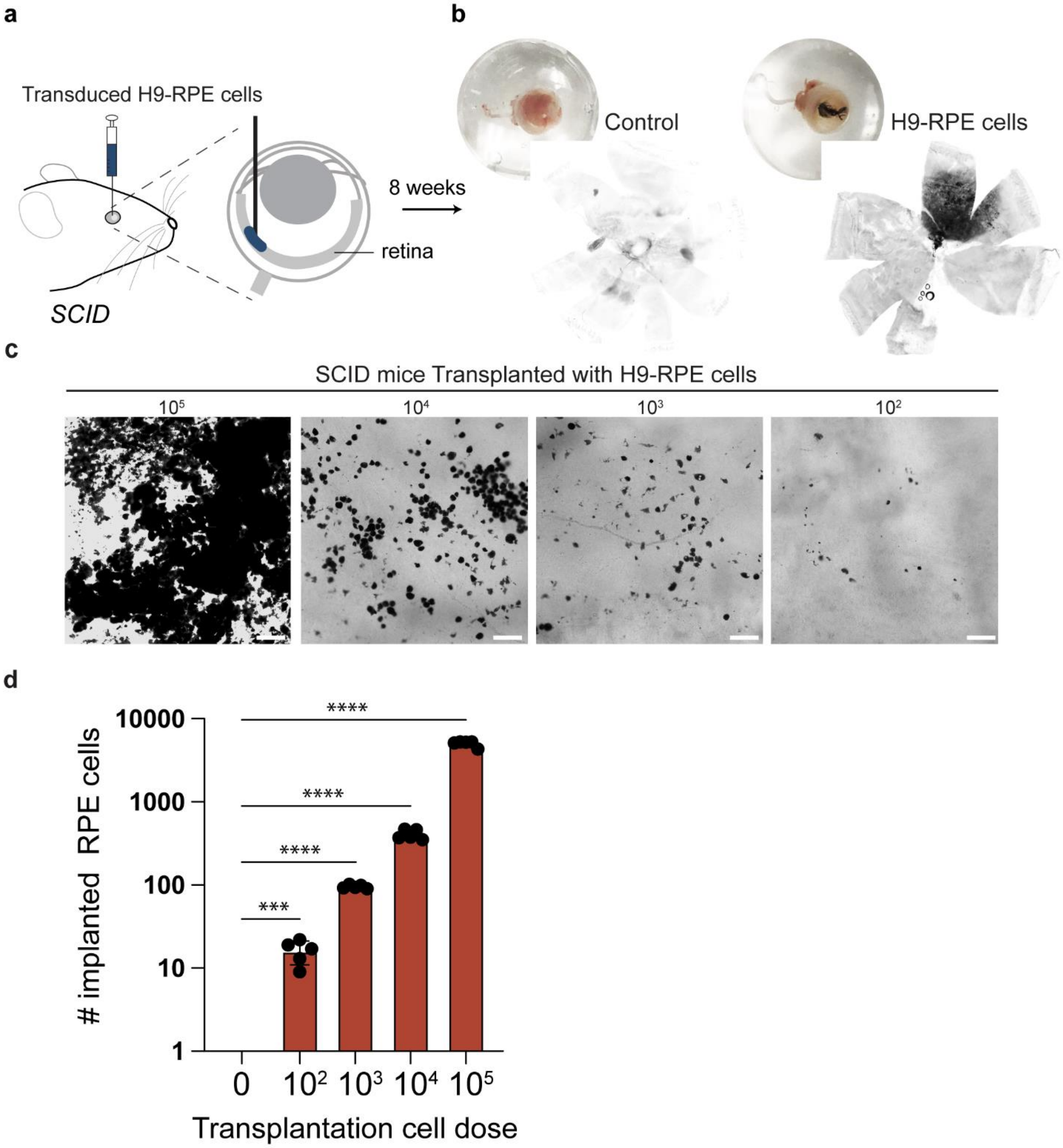


**Figure S4. Functional examination of SCID Retinas via RPE Transplantation. a.** Schematic showing transplantation of RPE cells into SCID mouse retinas. **b.** Representative images of eyes and retinal whole mounts from *SCID* control mice (no transplantation) and those that received $10^5$ RPE cells. **c.** Close-up images showing the viable transplanted RPE cells across varying injection doses. Scale, 20 µm. **d.** Quantification of the numbers of integrated RPE cells at 4 weeks post-transplantation. n=5 samples analyzed. Compared to the non-injected control, *Q*=0.000116 for $10^2$ cells implanted, and *Q*<0.0001 for the remaining groups (multiple *t-test* comparison with 5% FDR correction using Benjamini, Krieger, and Yekutieli method), ***$P<0.001$, ****$P<0.0001$. Data are shown as mean ± s.d.

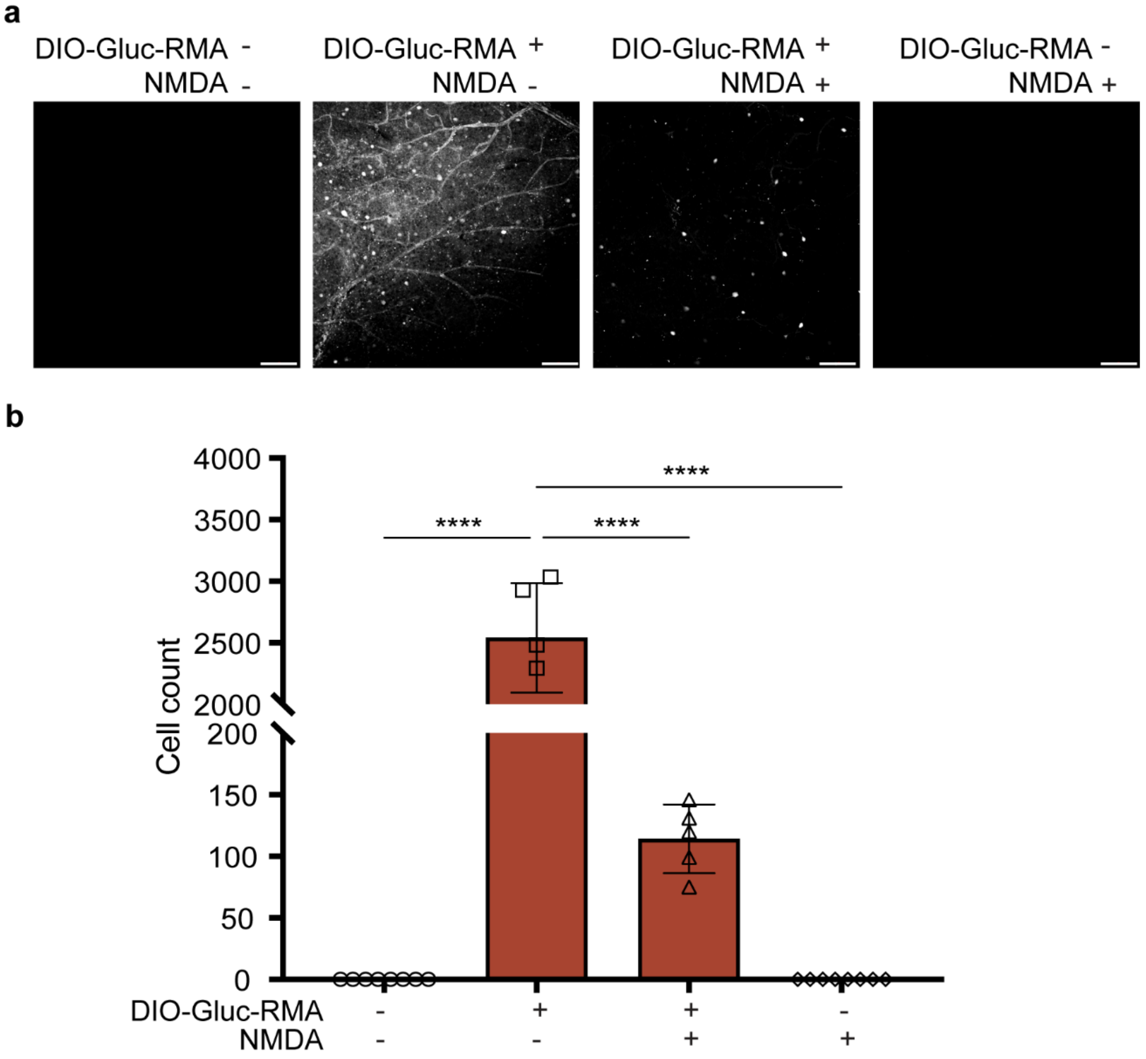


**Figure S5. Mouse RGC cell death after NMDA treatment. a.** Images showing the survival GlucRMA+ of after AAV-DIO-RMA with or without NMDA injections in *Vglut2Cr*e mice. Scale=100 µm. **b.** Quantification of the numbers of survival GlucRMA+ cells at 4 weeks post-AAV-DIO-RMA with or without NMDA injections. *n*=5 samples analyzed. Compared to the non-injected and NMDA-injected controls, *Q*<0.0001 for AAV-DIO-RMA and AAV-DIO-RMA+NMDA groups (multiple *t-test* comparison with 5% FDR correction using Benjamini, Krieger, and Yekutieli method), *****P*<0.0001. A reduction of

95.2% GlucRMA+ cells was detected in AAV-DIO-RMA+NMDA group compared to AAV-DIO-RMA. Data are shown as mean ± s.d.

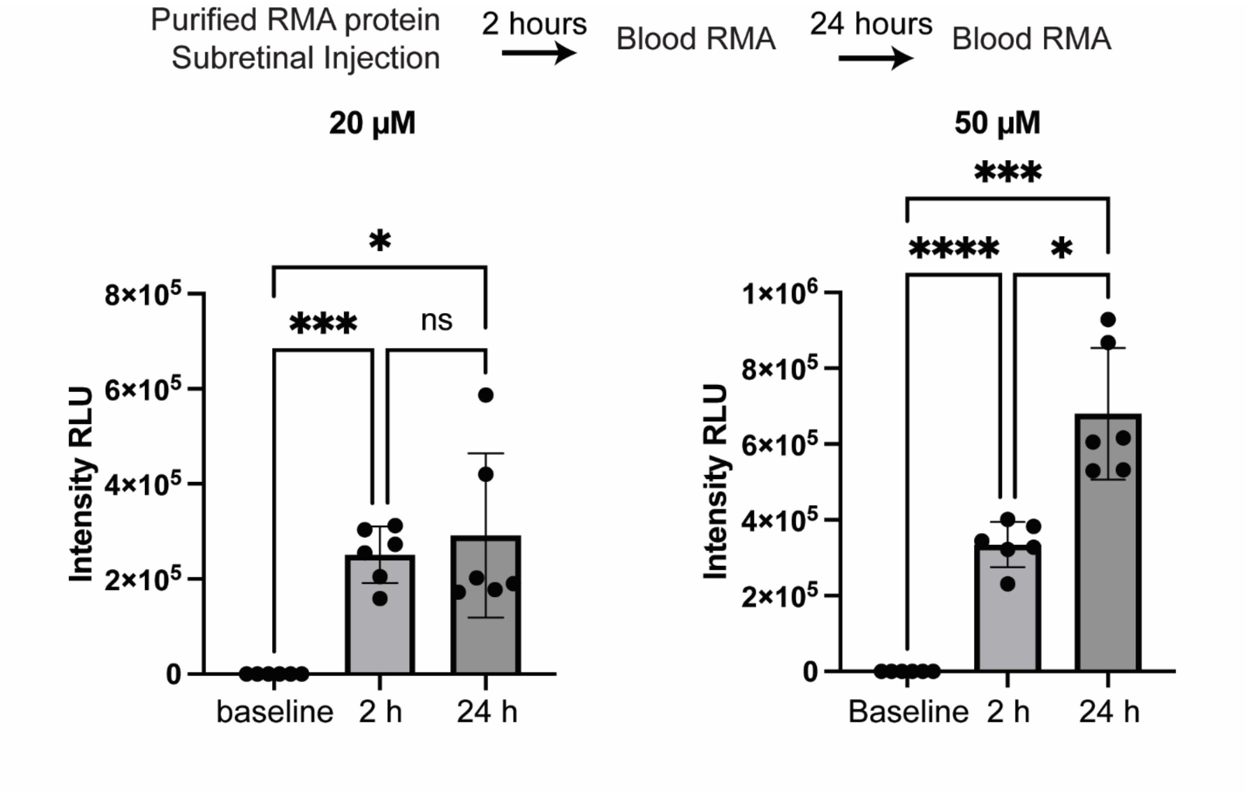


Figure S6. Blood RMA Levels Increase Following Administration of Purified GlucRMA Proteins. Quantification of blood RMA intensity increase after subretinal injections of 20 μM and 50 μM purified RMA proteins. Bloods were extracted before, 2 hours, and 24 hours after RMA protein injections. n=6 samples analyzed. Compared to the non-injected baseline controls, more than 10,000 folds elevations for both injected concentrations (multiple t-test comparison with 5% FDR correction using Benjamini, Krieger, and Yekutieli method), ****P<0.0001, ***P<0.001, and *P<0.05.
